# Challenges in estimating species age from phylogenetic trees

**DOI:** 10.1101/2023.11.19.567731

**Authors:** Carlos Calderón del Cid, Torsten Hauffe, Juan D. Carrillo, Rachel C. M. Warnock, Daniele Silvestro

## Abstract

**Aim:** Species age, the elapsed time since origination, can give an insight into how species longevity might influence eco-evolutionary dynamics and has been hypothesized to influence extinction risk. Traditionally, species ages have been measured in the fossil record. However, recently, numerous studies have attempted to estimate the ages of extant species from the branch lengths of time-calibrated phylogenies. This approach poses problems because phylogenetic trees contain direct information about species identity only at the tips and not along the branches. Here, we show that incomplete taxon sampling, extinction, and different assumptions about speciation modes can significantly alter the relationship between true species age and phylogenetic branch lengths, leading to high error rates. We found that these biases can lead to erroneous interpretations of eco-evolutionary patterns derived from the comparison between phylogenetic age and other traits, such as extinction risk.

**Innovation:** For bifurcating speciation, which is the default assumption in most analyses, we propose a probabilistic approach to improve the estimation of species ages, based on the properties of a birth-death process. We show that our model can reduce the error by one order of magnitude under cases of high extinction and high percentage of unsampled extant species.

**Main conclusion:** Our results call for caution in interpreting the relationship between phylogenetic ages and eco-evolutionary traits, as this can lead to biased and erroneous conclusions. We show that, under the assumption of bifurcate, it is possible to obtain better approximations of species age by combining information from branch lengths with the expectations of a birth-death process.

## Introduction

The estimation of species age, or the elapsed time since species origin, is important to evaluate mechanisms that link species longevity with eco-evolutionary processes (Benton, 2013; Swenson, 2019). For instance, age-dependent extinction hypotheses test the relationship between species age and extinction probability, assessing whether extinction rates differ between young and old species (Balmford, 1996; Eldredge et al., 2005; Pearson, 1995). Likewise, species age could be a measure of colonization time, especially in island systems (Tanentzap *et al*. 2015) or during biotic invasions triggered by geological events, such as the formation of the Central American Isthmus for the Great American Interchange (Carrillo *et al*. 2015, 2020). Species age is measured in the fossil record through different statistical and probabilistic approaches, based mostly on taxa’s stratigraphic duration (i.e., the time between the first and last appearance of a taxon in the fossil record) (Foote, 1996; Foote & Raup, 1996). Several of these approaches consider differences in fossil sampling and temporal resolution (Alroy et al., 2001; Silvestro et al., 2019). Species ages estimated from paleobiological data offer a reliable measure of species’ temporal duration which can be used in macroevolutionary studies (Benton, 2016; Silvestro et al., 2020; Van Valen, 1973). More recently, several studies have used the length of terminal branches in time-calibrated phylogenies as a proxy for the age of extant species, an approximation that we hereafter refer to as “phylogenetic age” (Alzate et al., 2023; Davies et al., 2011; Gaston & Blackburn, 1997; Johnson et al., 2002; Pie & Caron, 2023; Sonne et al., 2022; Tanentzap et al., 2020; Verde Arregoitia et al., 2013). These phylogenetic ages have been used as the basis to test for links between species age and current extinction risks (Tanentzap et al., 2020; Verde Arregoitia et al., 2013) and to assess various correlations with evolutionary, biogeographical, and ecological patterns in living species (Alzate et al., 2023; Freer et al., 2022; Kennedy et al., 2022; Pie & Caron, 2023)

While several studies have used phylogenetic age at face value for species age (e.g., Johnson et al. 2002; Tanentzap et al. 2020; Verde Arregoitia et al. 2013), their potential deviation from the true species ages remains unclear. Specifically, we identify three non-mutually exclusive shortfalls that can lead to over- or underestimation of species ages. First, incomplete sampling of extant species, either due to incomplete species sampling or linked to species still being unknown to science, can bias phylogenetic age estimation by artificially increasing the length of terminal branches (Heath et al., 2008; Mynard et al., 2023).

Second, extinction events will mask branching events in phylogenetic trees of extant species (Harvey et al., 1994; Nee & May, 1997). Even in phylogenetic trees that include extinct taxa, the incompleteness of the fossil record will inevitably lead to missing lineages and incorrect topologies. Unsampled extant and extinct species from the phylogeny results in an inflation of the length of terminal branches leading to sampled species (i.e., the tips of the tree), thus altering phylogenetic species ages. For instance, if the extinct species of the *Homo* genus are not included in a phylogeny, the phylogenetic age of *Homo sapiens* is approximately 10 million years, i.e., the age of the last common ancestor with its sister species, the chimpanzee (Rivas-Gonzáles et al. 2023). This estimate exceeds the age of the oldest known fossil of modern humans (i.e., *Homo sapiens*) by two orders of magnitude (Fig. 1; Callaway 2017).

**Fig. 1.**
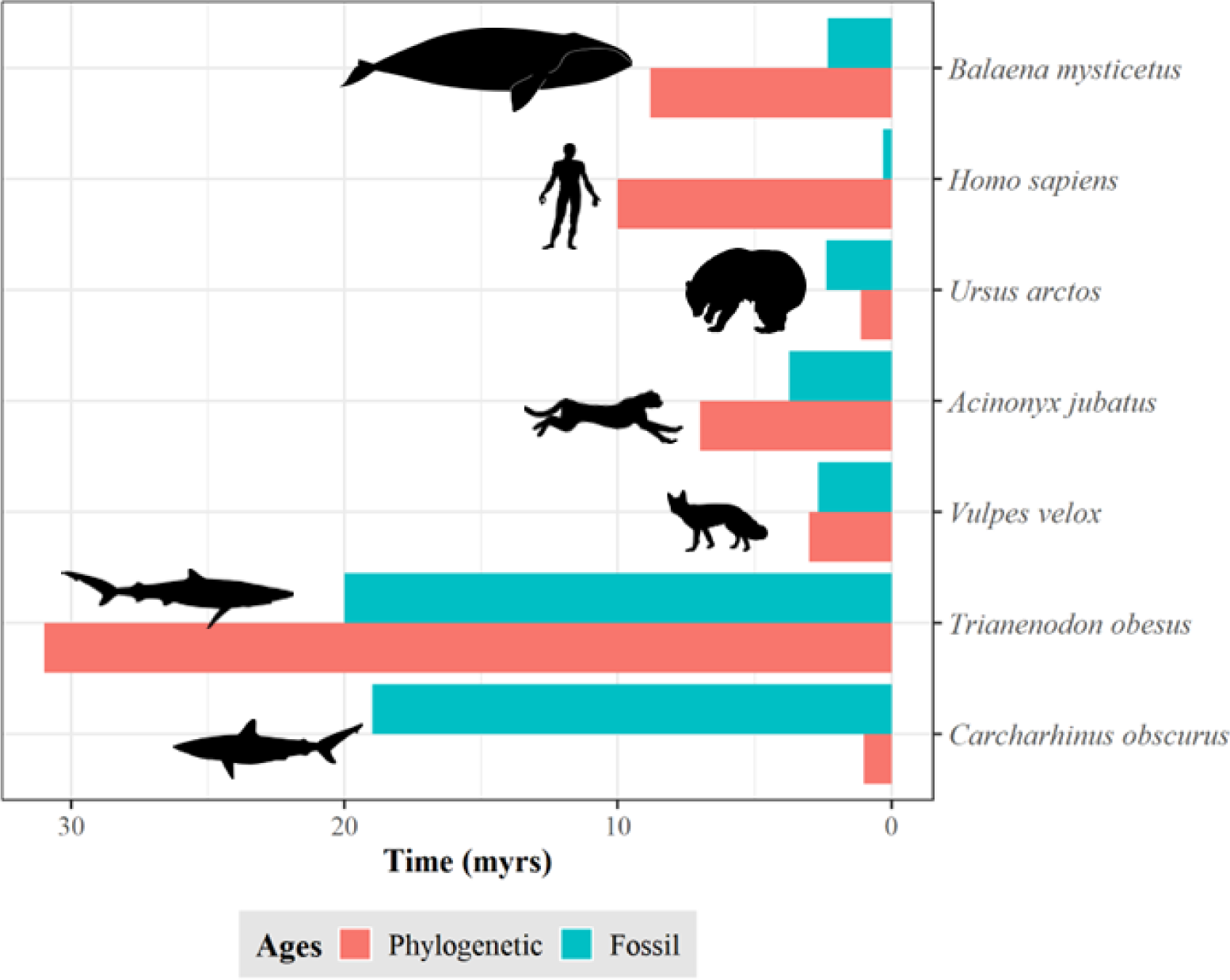
Discrepancy between species ages. Estimations based on the length of the terminal branch in a time-calibrated phylogeny (red) and the first appearance in the fossil record (green). Shark data (*Trianenodon obesus* and *Carcharhinus obscurus*) obtained from Brée et al. (2022). Mammals’ phylogenetic data (except *Homo sapiens*) obtained from Upham et al. (2019). Mammals’ fossil data (except *Homo sapiens*) obtained from Silvestro *et al*. (2018). *Homo sapiens* fossil and phylogenetic data obtained from Callaway (2017) and Rivas-Gonzáles et al. (2023), respectively.

The third shortfall, is that the tree alone does not contain information about the underlying speciation mode and does not include species labels along its branches, such that only the tips can be unequivocally assigned to a named species (Losos & Glor, 2003). Alternative speciation modes have been discussed in the literature reflecting different biological processes and species concepts, including bifurcating, budding, and anagenetic speciation (Foote, 1996; Silvestro et al., 2018). These modes define the relationship between the ancestral species and its descendants, thus contributing to determining species ages (Rosenblum et al., 2012; Wagner, Erwin, & Anstey, 1995) (Fig. 2). Most phylogenetic trees are depicted in the rectangular shape, where the two descending lineages split symmetrically from an ancestral lineage, thus suggesting a bifurcating speciation mode where two new species replace the ancestral lineage (Baum et al., 2005; Caetano & Quental, 2022). However, the often-unstated assumption of virtually all birth-death processes used to model phylogenetic branching times, is that speciation occurs as a budding process, with a speciation event leading to a single new species and the survival of the parent species, even though we cannot determine which descendant branch is the new species (Gernhard, 2008; Nee et al., 1994; Stadler, 2013; but see Stadler et al., 2018). Anagenetic speciation, in contrast, does not lead to a branching event and is therefore not ordinarily visible on a phylogenetic tree.

**Fig. 2.**
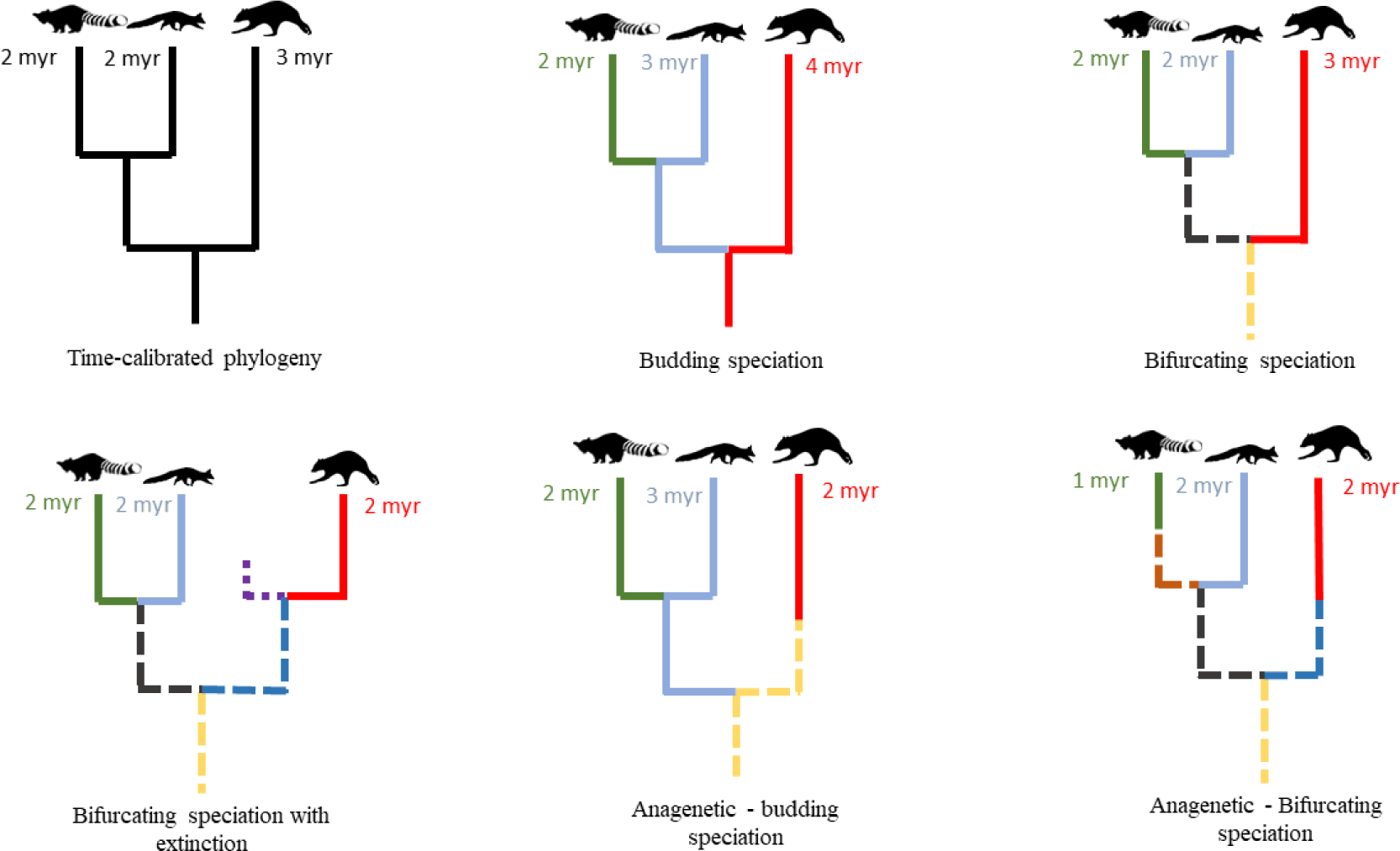
Impact of speciation mode and extinction on species age. For the same hypothetical time-calibrated phylogeny of extant species, the continuation of the same color indicates the same species, solid and dashed lines represent extant and extinct species, respectively, under different speciation modes and extinction scenarios. Numbers display the resulting age of the respective species in million years (myr).

All speciation modes may reflect plausible biological processes, and evidence for each mode has been found in the fossil record (Foote, 1996; Simpson, 1984) and in extant species (Skeels & Cardillo, 2019). Under bifurcating speciation mode, phylogenetic ages equal species ages when there is no extinction, and all species are sampled. A phylogenetic tree, under budding speciation, typically lacks information about which of the two descendent branches is the new species after a speciation event (but see Aze *et al*. 2011). Thus, in the absence of extinction, the phylogenetic age of one sister species will equal its species age while the other will be older but without the possibility to identify which one is which. Anagenetic speciation is not associated with a branching event but can be used to describe different species or morphospecies (Emerson & Patiño, 2018) delimited by substantial phenotypic change occurring along a lineage (Roopnarine et al., 1999) and will cause a higher phylogenetic age than the genuine species age.

Here we use simulations to quantify the predictability of species age from phylogenetic trees of extant taxa, under different diversification scenarios. Specifically, we performed simulations where we know the true age of species to: 1) quantify the error in phylogenetic ages under various scenarios combining different speciation modes with a range of speciation and extinction rates and incomplete sampling; 2) examine whether this error affects our ability to make qualitative age comparisons between species; 3) explore whether the signal of age-correlated extinction risk is preserved in the phylogenetic age of species. Finally, we propose a new method to estimate species age more accurately under the assumption of bifurcating speciation, while correcting for incomplete species sampling,, and assess its ability to improve our interpretation of age-dependent extinction risks.

## Methods

### Simulating species ages

We generated complete phylogenies of extant and extinct species under a stochastic birth-death process using the package TreeSim 2.4 (Stadler, 2010) for the R 4.3.0 statistical programing environment (R Core Team 2023). Then we mapped species on the complete phylogenies using the R package FossilSim 2.3.1 (Barido-Sottani et al., 2019) under different speciation modes, thus assigning species labels across all branches of the tree. We used the labels assigned to terminal extant taxa to determine the true species ages. We then dropped all extinct species from the tree and obtained the length of terminal branches, to quantify the phylogenetic age of extant species. Finally, we rescaled all phylogenetic trees to a root age of one, which ensures that the absolute errors in species ages are comparable in plots, and compared the relative true and phylogenetic ages among different simulation scenarios.

### Error in equating phylogenetic and species age

To explore whether there is a consistent over- or underestimation of species ages and to quantify error in approximating species ages with phylogenetic ages, we simulated a range of datasets with different speciation modes and diversification rates. First, we simulated 3 sets of 100 phylogenetic trees with 100 extant species based on speciation rates equal to 0.1, 0.5, and 1, combined with 100 extinction rates ranging from 0 to 0.99 in equal increments (Beaulieu & O’Meara, 2016). Second, on each of these phylogenies we mapped species according to different scenarios of speciation: (1) budding speciation, (2) bifurcating speciation, (3) a combination of budding speciation and anagenetic speciation with the rate of anagenesis set to half of the speciation rate, and (4) bifurcating speciation combined with anagenetic speciation with the rate of anagenesis set to half of the speciation rate.

Across all trees, we obtained in total 120,000 extant species, 30,000 for each speciation scenario. For each speciation mode and extinction fraction rate (defined as death/birth Beaulieu & O’Meara, 2016), we calculated the mean absolute percentage error (MAPE) across all species for each tree as measure of the deviation between the phylogenetic ages from the true age:

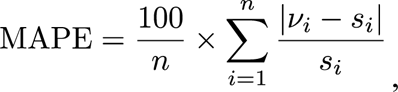

where *s* is the true species age, *v* is the phylogenetic age, and *n* is the number of tips in the tree (pruned of the extinct species).

Next, for each speciation mode, we plotted the MAPE against the simulated extinction fraction.

### Impact of age error in comparing species ages

To explore whether the error introduced by approximating species age with phylogenetic age impacts our ability to make qualitative judgements, such as which of two extant species is the younger one, we simulated 1,000 phylogenetic trees with values of extinction fractions (extinction rate divided by speciation rate) of 0.9, 0.5, and 0, combined with a fixed speciation rate of 1 (3000 trees). Second, on each of these phylogenies, we mapped species according to budding and bifurcating speciation. Thus, we simulated 300,000 extant species for each speciation mode. Next, we calculated the proportion of cases where the younger of two species, according to its phylogenetic age, is, in fact, the older one given the true age of the two species. We performed this comparison from the perspective of an empirical researcher that can only obtain the phylogenetic ages. We made two types of comparisons for each phylogeny: (1) between the youngest and oldest species in the phylogeny, and (2) between two randomly selected species.

### Error in the phylogenetic age due to uniform incomplete sampling

To evaluate the error of equating phylogenetic and true age that is introduced by uniform incomplete sampling under bifurcating speciation, we simulated 1,000 phylogenetic trees with a speciation rate of 0.3 and extinction rates of 0.05, 0.15, and 0.25.. In addition to fully sampled phylogenies (where all extant species are included), we also simulated trees with incomplete taxon sampling, where 25% or 50% of the tips were randomly removed. We simulated trees conditioning on the number of tips, such that they included 100 sampled tips even after dropping the unsampled ones (134 or 200 total tips). We calculated the MAPE for each tree and compared the incomplete sampling percentages for each extinction scenario.

### A probabilistic method to infer species age

Under the assumption of bifurcating speciation, the phylogenetic age represents the upper boundary of plausible species ages and corresponds to the true age in the absence of extinction and complete taxon sampling. However, the true age could be younger if extinction led to the disappearance of recent cladogenetic events from the phylogeny of extant species and/or if incomplete sampling led to unobserved branching events. Given a phylogenic age *v_i_* the probability that the true species age *s_i_* is exactly *v_i_* is conditional on no other speciation or extinction event having occurred between *v_i_* and the present. We approximate the probability of no speciation or extinction for an arbitrary small time bin *t* based on the probability that a lineage results in a single descendant, which is (Kendall, 1946):

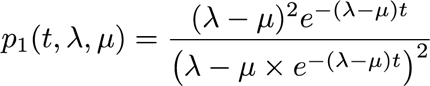

based on a birth-death process with time-homogenous speciation rate λ and extinction rate μ. The probability that no event occurs over a time window *v, i.e.,* until the age of the observed node *v_i_*, is approximated as:

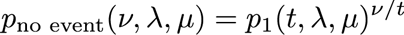

In the absence of extinction (μ = 0) and full taxon sampling, the probability of the true species age *s_i_* to be equal to *v_i_* is 1, because any speciation event following the node *v_i_* would be observed in the tree of extant species. This probability is however further incomplete sampling. Thus, we calculate the normalized probability of the speciation event to occur at time *v_i_* as:

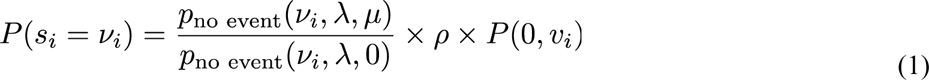

where ρ is the probability of sampling lineage *i*, i.e. the sampling fraction (ρ = 1 if all living species are included in the tree) and *P(0, v)* is the probability that the sister lineage leaves at least one sampled descendant (without which the node *v_i_* would not be observed), as defined by Yang and Rannala (1997):

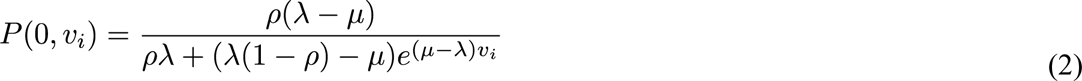

We then compute the probability of a speciation time for any given time *τ* as:

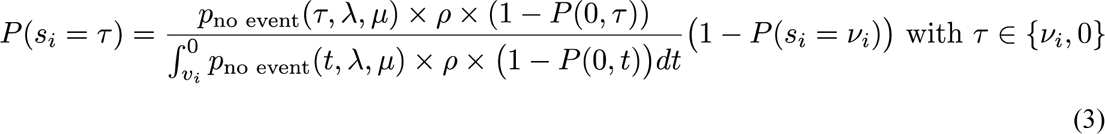

where the first term quantifies the probability of a branching event giving origin to species *i* after which no other speciation or extinction events occur along the lineage between time τ and the present. We note that the probability includes the sampling probability of species *i* (ρ) and the probability that the sister lineage originated at time τdid not leave any sampled descendants (1 – P(0, τ)). The second term normalizes it after accounting for the probability that speciation occurred exactly at the node *v_i_*. We use equations (1) and (3) to approximate a density describing the probability of a species origination at the observed phylogenetic age and along the branch connecting the node with the tip. We tested the mean and the median of the distribution as point estimates of species age compared them to the phylogenetic age.

To evaluate the accuracy of our probabilistic species age estimator, we applied it on the simulations described in the previous section and calculated the MAPE for each tree as a measure of the deviation between the function’s point estimates and the true age. Then, we calculated the ΔMAPE as the difference between the MAPE of our function’s point estimates and the MAPE of the corresponding phylogenetic ages.

### Simulation of age-dependent species extinction risks

To evaluate the impact of the erroneous estimation of species age due to incomplete sampling and extinction on macroevolutionary analyses, we explored whether a relationship between species age and contemporary extinction risk (e.g., Johnson et al. 2002; Tanentzap et al. 2020; Verde Arregoitia et al. 2013) is correctly preserved in the phylogenetic ages. For this, we binned the same number of extant species according to their age and assigned them to five categories simulating an increase in extinction risks with age encapsulated by the IUCN categories: Least Concern (LC), Near Threatened (NT), Vulnerable (VU), Endangered (EN), and Critically Endangered (CR; International Union for the Conservation of Nature 2016). With this, we generated a positive effect with older species being at higher extinction risk regarding the IUCN categories, assuming bifurcating speciation.

Then, we quantified the share of the 1000 datasets where the order of the mean age per IUCN category did not match with the simulated monotonic increase when utilizing (a) phylogenetic ages, and (b) the probabilistic species age estimator. We also evaluated the effect of nonrandom incomplete sampling (older species were less prone to be sampled than younger ones) on the evaluation of species age-correlated extinction risks and the ability of our probabilistic function to reduce the error rates.

## Results

### Error in equating phylogenetic and species age

Under the assumption of bifurcating speciation and with no extinction events, phylogenetic ages matched the true age of extant species (Fig. 3). At low extinction fractions (< 0.25), 96% of the phylogenetic age estimations were congruent with the true age. At higher extinction fractions (> 0.75), this was also the case for most species (73%). However, age overestimation increased with extinction fraction and in some cases the phylogenetic age erroneously suggested that the species is as old as the root age. While under bifurcating speciation, the phylogenetic age never underestimated the true species age, both over- and underestimation occurred in the case of budding speciation. Moreover, the proportion of cases where the phylogenetic ages equal the species age was lower than in the bifurcating scenario (Fig. 3). Overestimated ages were more frequent with high extinction while underestimations occurred with low extinction, but in principle both happened under the complete range of extinction rates (Fig. 3). Even at low extinction fractions, ∼50% of phylogenetic ages did not match the true ages.

**Fig. 3.**
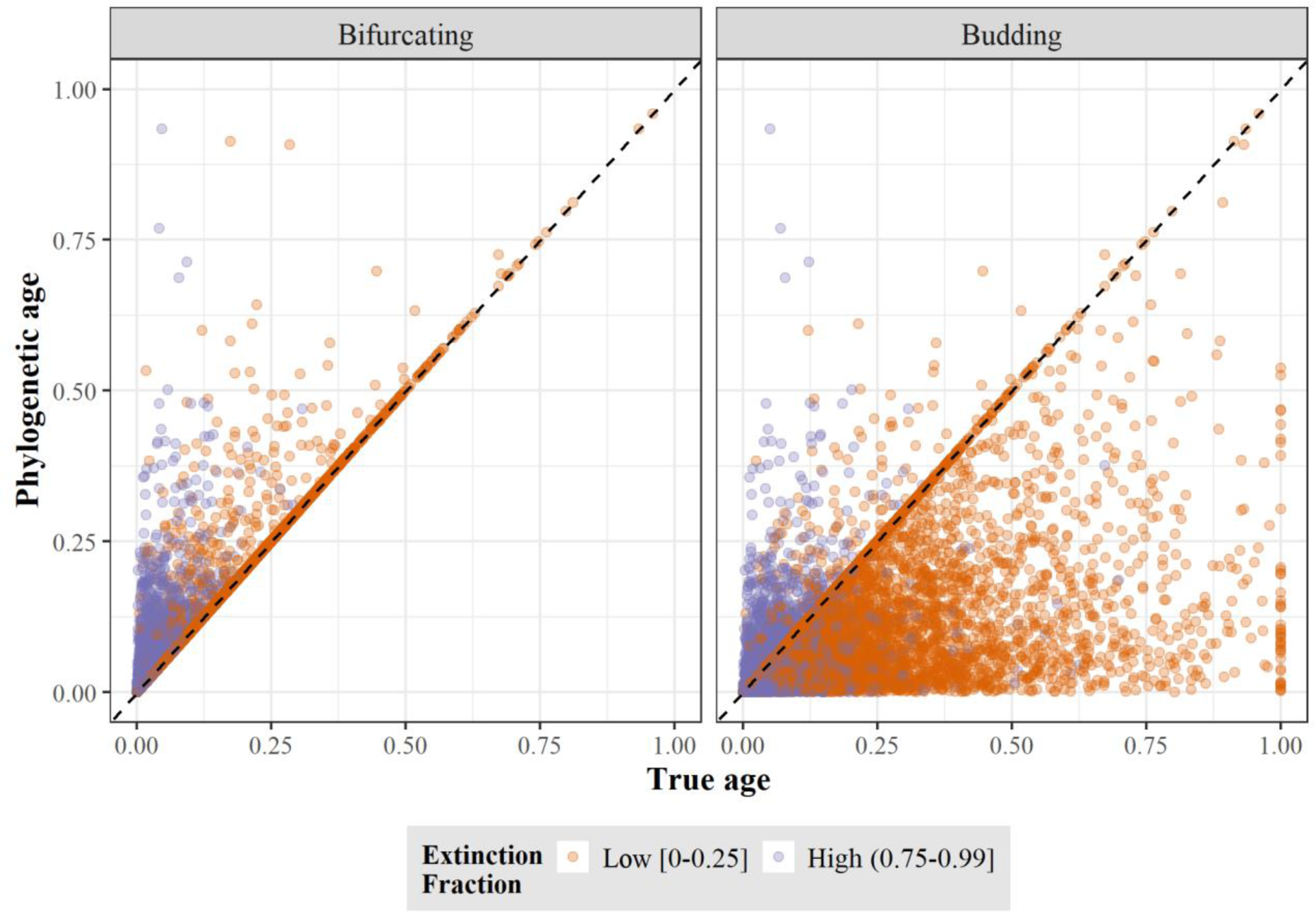
True age versus phylogenetic age at low and high extinction fraction for bifurcating and budding speciation. Each point represents a species and both ages, true and phylogenetic, are scaled to the root age of the correspondent phylogenetic tree.

In datasets simulated under a mixture of anagenetic and bifurcating speciation, phylogenetic ages deviated more strongly from the true ages than under a pure bifurcating process, given that anagenetic events are unobserved in the phylogeny (Fig. SM1). With a low extinction fraction, phylogenetic ages were congruent with the true species ages in 78% of the cases instead of 96%, and with high extinction the share decreases from 73% to 62%. Datasets with mixed anagenetic and budding speciation, phylogenetic ages also deviated more that under a pure budding process; with a low extinction fraction, phylogenetic ages were congruent 40% instead of 50% of the time, and with high extinction the accuracy decreased to 34%.

While a budding speciation mode led to a higher baseline error than bifurcation, the latter showed a stronger increase with extinction (Fig. 4). Under both modes of speciation, speciation rates did not have a substantial impact on error in age. For strictly bifurcating speciation, there was no error in the absence of extinction, but the MAPE increased to up to 150% with extinction fractions exceeding 0.75. In contrast, under budding speciation the MAPE was around 25% in the absence of extinction, increasing to 30-120% with extinction fractions exceeding 0.75. In datasets that included anagenetic speciation, the MAPE reached as high as 500% in some simulations (Fig. SM2).

**Fig. 4.**
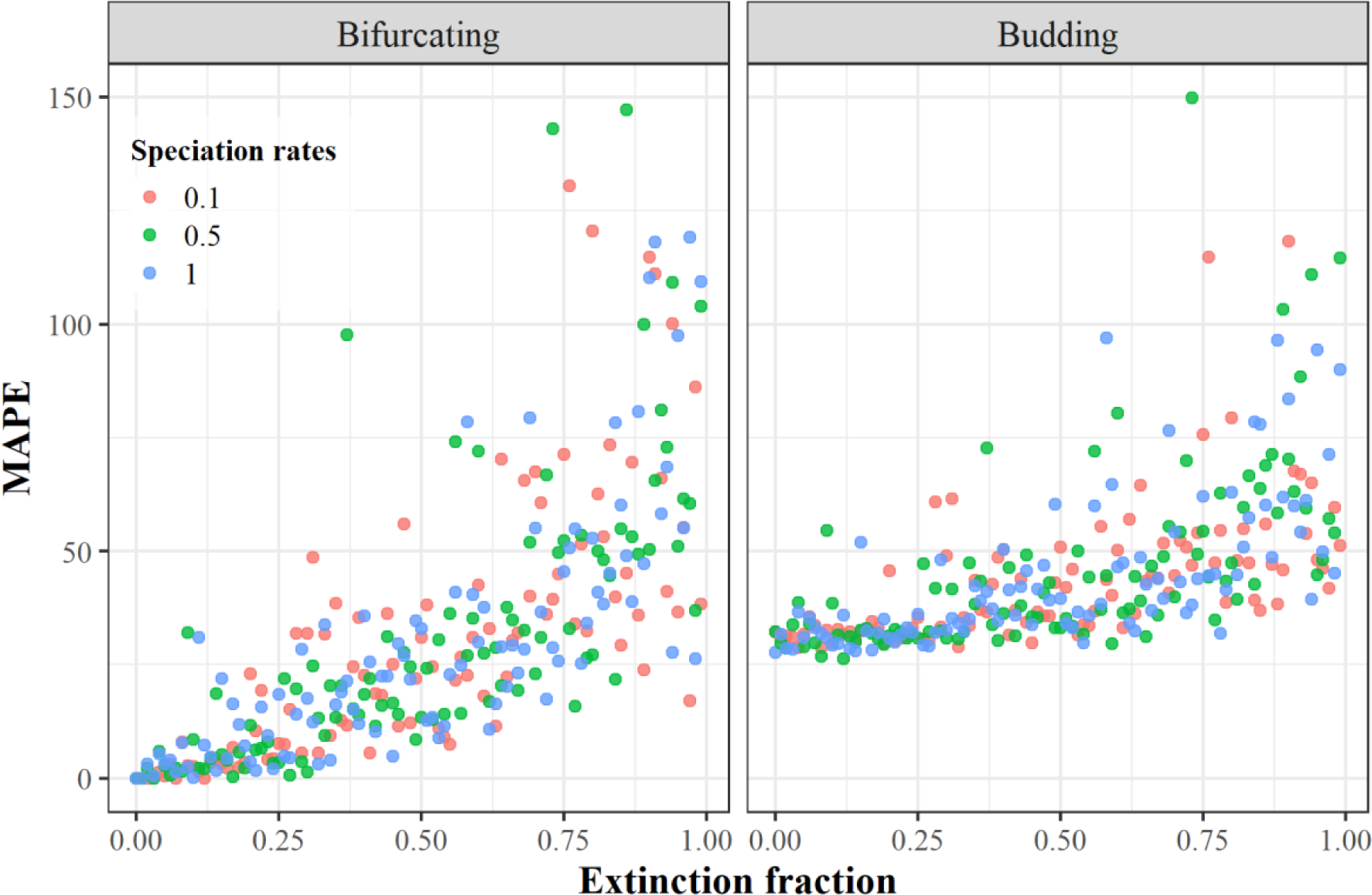
Error in equating phylogenetic age with true age. The error was quantified as mean absolute percentage error (MAPE) between the true and phylogenetic ages across all species for each tree simulated under bifurcating and budding speciation. Each dot represents one replicate of the 300 trees for each speciation mode using different rates of speciation and extinction fraction.

### Impact of age error on comparing species ages

For the combination of strictly bifurcating speciation and all extinction scenarios, selecting the phylogenetically youngest and oldest species never resulted in a case where the presumed older species has been in fact the younger of the two species according to their simulated age (Fig. SM3a). Thus, for this speciation mode, the risk of a qualitative error when comparing species at the extremes of the age range is minimal. However, the age ranking of two randomly selected species was found to be incorrect in 6% and 8%, for intermediate and high extinction, respectively (Fig. SM4a). Thus, qualitative errors in comparing species ages are non-negligible under the assumption of bifurcating speciation.

In contrast, for budding speciation, the age rank of the oldest and youngest species was erroneously determined in 2.2% of the simulations in the absence of extinction, increasing to 7.5% and 12.2% for intermediate and high extinction, respectively (Fig. SM3b). Thus, under the assumption of budding speciation, there is a substantial risk of mistaking the oldest and youngest species in the clade. The error in age ranking of two randomly selected species was even higher, exceeding 25%, irrespective of the extinction level (Fig. SM4b).

### Error on equating phylogenetic and species age given uniform incomplete sampling

In the absence of extinction and assuming bifurcating speciation, the MAPE for fully sampled trees was ∼ 0 %, but it increased to ∼100% and ∼300% for 25% missing species and 50% missing species, respectively (Fig. 5). With intermediate extinction, the MAPE for fully sampled trees was 25 ± 20%, but increased 15-fold for trees missing 25% of the extant species; and for trees missing 50 % of the extant species the error increased 85-fold. Under high extinction, the MAPE for fully sampled trees was 60 ± 38%; for trees missing 25% of extant species the error increased 8-fold of magnitude; and for trees missing 50% of extant species the error increased 23-fold.

**Fig. 5.**
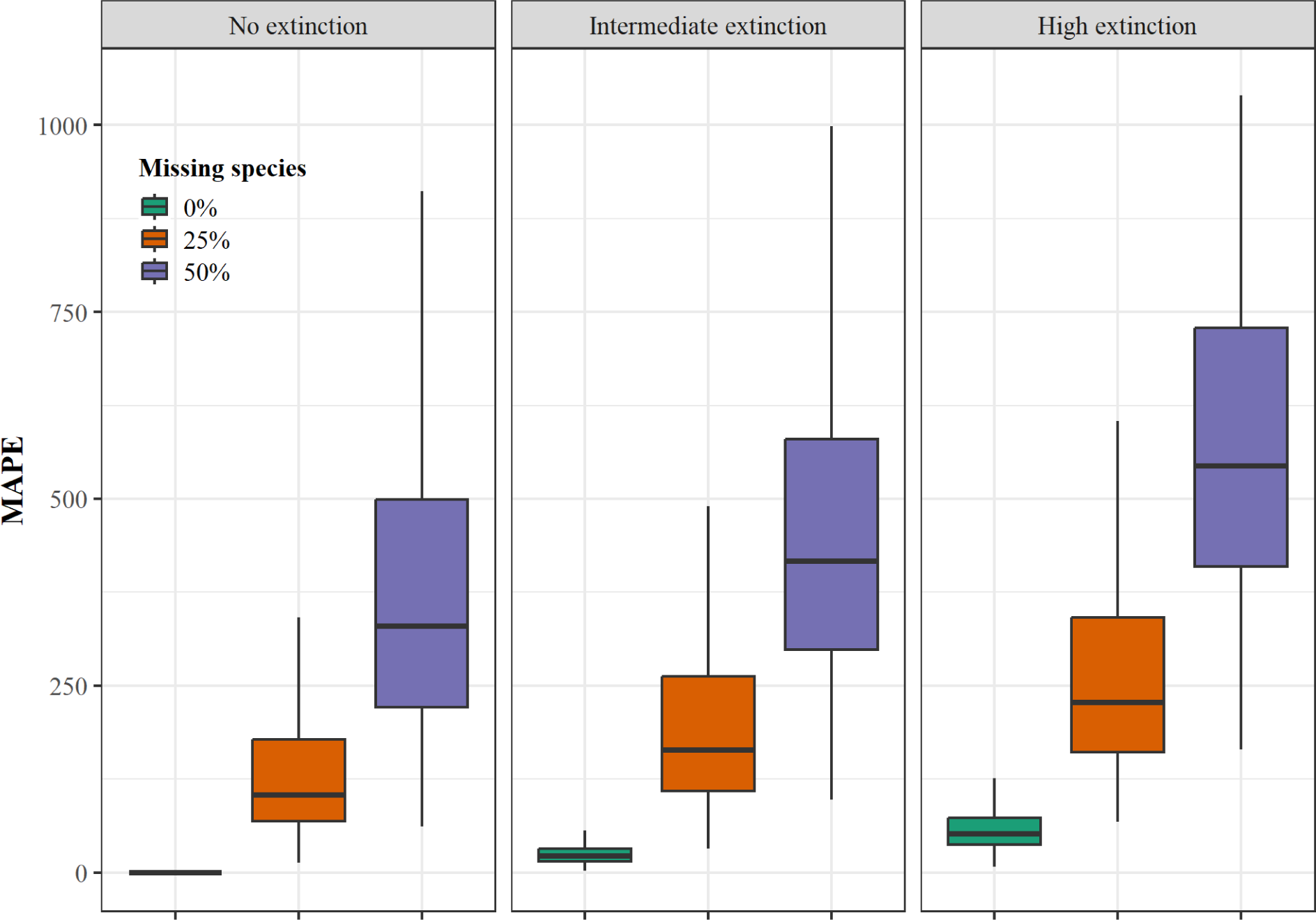
Effect of incomplete taxon sampling and extinction on error in species ages. Error in equating the phylogenetic age with true species age for the fully sampled phylogeny, and 25% and 50% of missing extant species, in the three extinction scenarios (no extinction, intermediate, and high; from left to right). The error was quantified as mean absolute percentage error (MAPE) between the true phylogenetic ages across 100 species for each of 1000 trees for each missing species scenario simulated under bifurcating speciation.

### Probabilistic species age estimation

With increasing extinction and missing species, our probabilistic estimation of species ages resulted in a substantially lower error compared with the phylogenetic age (Figs. 6, SM5). Under low extinction and fully sampled trees the MAPE was slightly worse compared to the use of phylogenetic ages (ΔMAPE = 0.59 ± 1.1 % when using the mean of the estimated ages and 2.1 ± 1.3 % for the median across estimates). While with high extinction and a taxon sampling of 50% species the probabilistic estimations reduced the error by 2 orders of magnitude regarding the phylogenetic age (ΔMAPE = −476% for the mean estimated ages and −507 % for the median across estimates; Figs. 6, SM5).

**Fig. 6.**
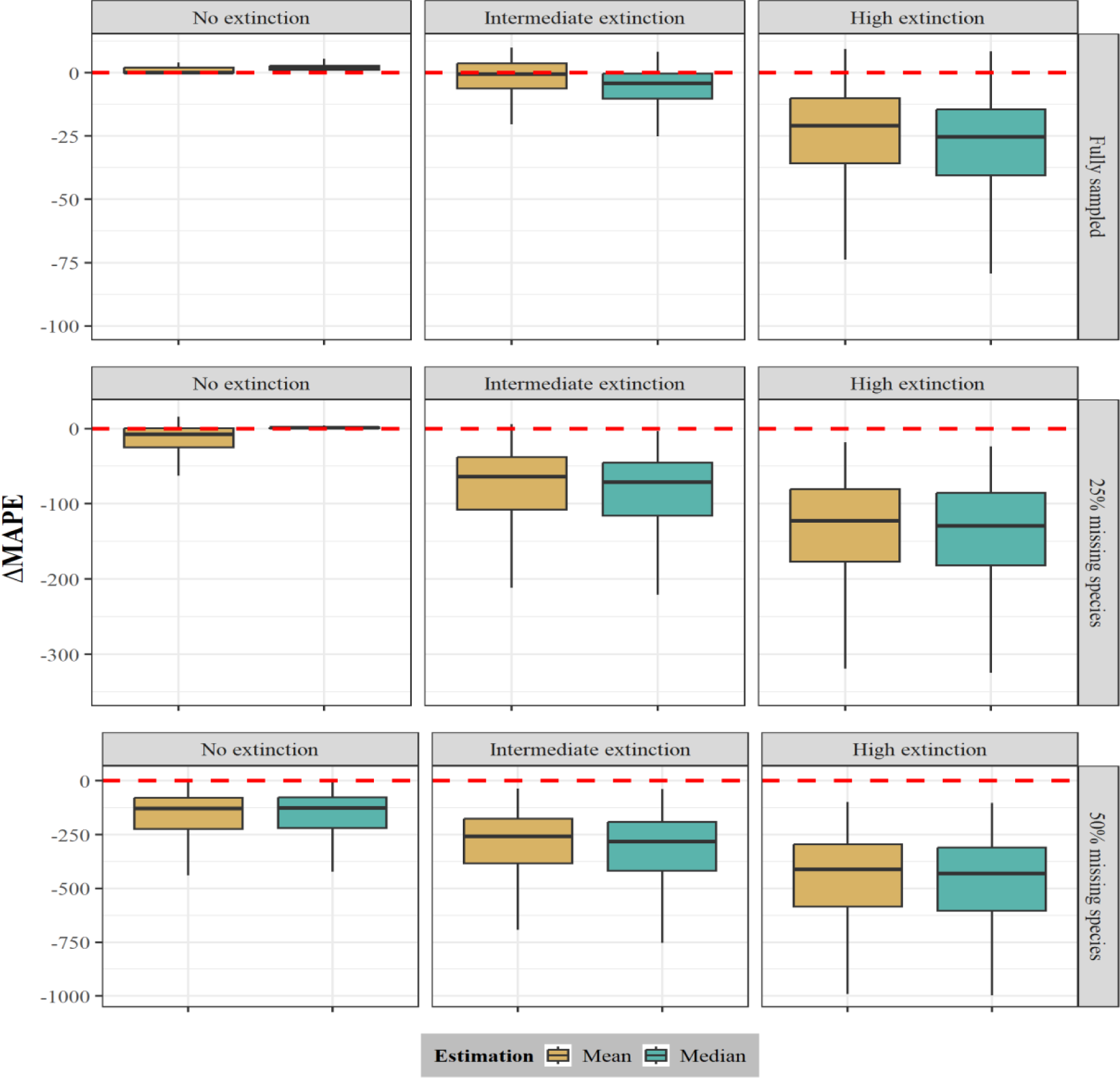
Performance of the probabilistic age estimator. ΔMAPE for the three extinction scenarios (low, intermediate, and high; from left to right) and the three sampling scenarios (full, 25%, and 50% missing species; from up to down) was quantified as the difference of the mean absolute percentage error (MAPE) of the probabilistic estimator point estimates (mean and median) and the MAPE of the phylogenetic age. The MAPE was quantified as the difference between the true and point estimates (mean and median) of phylogenetic ages for 100 species across 1000 trees for each extinction scenario simulated under bifurcating speciation. The red dashed line represents no difference between the compared MAPEs, negative ΔMAPE values indicate an improvement in the accuracy of the probabilistic estimator over the phylogenetic age. We note that for clarity, we used different scales for the Y-axis for each sampling fraction.

### Detecting age-dependent extinction risk

The use of phylogenetic age as an approximation of species age led to error rates of 1.3, 7.2, and 18.6 % in detecting the correct monotonic correlation between species ages and extinction risk, with fully sampled trees, and for scenarios with low, intermediate, and high extinction rates, respectively (Fig. SM6). Thus, even under intermediate extinction the true relationship between age and extinction risk was wrongly estimated in a significant fraction of the simulations, and higher extinction rates led to a further substantial drop in the reliability of this approach. In contrast, estimating species ages based on our probabilistic method led to much lower error rates (3 to 4-fold) that dropped to 1.4% and 3.1% with intermediate and high extinction, respectively. Under incomplete taxon sampling, in which the sampling probability was negatively correlated with species age, and under intermediate extinction, the error rates increased to 13.1%, and 51.9% for scenarios with 25%, and 50% of missing extant species, respectively (Fig.7). For 25% missing species, the function’s point estimates reduced the error rates by 4 to 7 orders of magnitude, decreasing to 3.1% for mean age and 1.7% for median age. For 50% missing species, the point estimates reduced the error rates (2.5 to 3.7-fold), decreasing to 21.1% for mean age and 14.2% for median age.

**Fig. 7.**
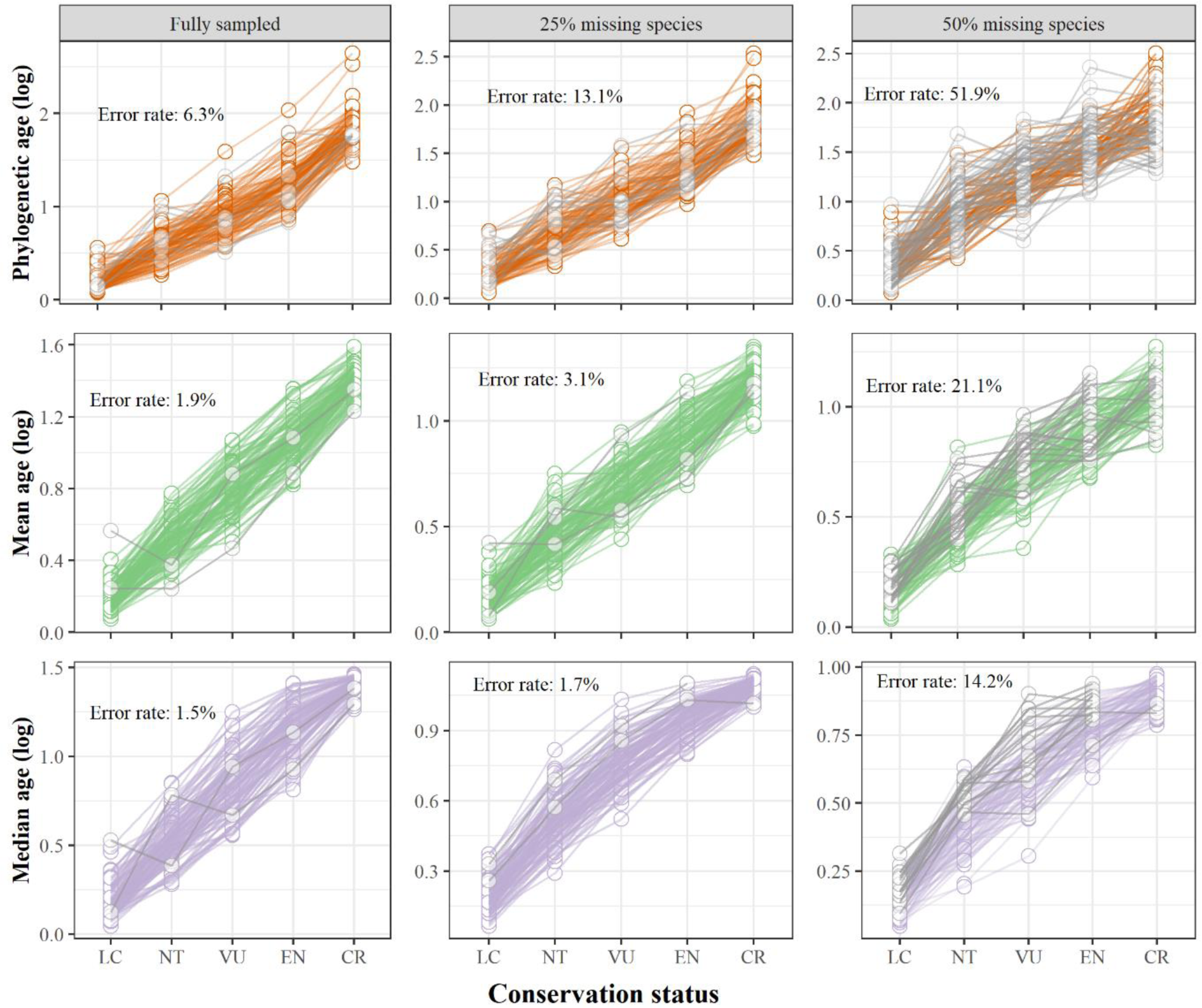
Power to recover an age extinction-risk relationship under different incomplete sampling scenarios. Simulated species ages under an intermediate extinction scenario and assuming bifurcating speciation were binned into conservation status categories, which represents an increase in extinction risk by true age (LC = Least Concern; NT = Near Threatened; VU = Vulnerable; EN = Endangered; CR = Critically Endangered). We used the phylogenetic age (orange), the mean age (green), and median age (purple) obtained from our function’s point estimates to calculate the mean age per conservation status category and assess if every mean age increases in comparison with the previous category with lower extinction risk. We evaluated this relationship in three sampling scenarios (fully sampled, 25%, and 50% of missing species; from left to right). The error rate is the percentage across all 1000 phylogenies where the relationship between the mean ages and the conservation status categories is not always increasing (shown by grey lines).

## Discussion

The use of branch lengths of phylogenetic trees as an approximation of species ages is becoming central to an increasing number of studies that use them to evaluate the relationship with macroecological or evolutionary patterns, for instance current extinction risks (Pie & Caron, 2023; Sonne et al., 2022; Tanentzap et al., 2020; Verde Arregoitia et al., 2013). Here we showed that this approximation leads to substantial errors and that its accuracy is hampered by three shortfalls: incomplete sampling of extant species, unobserved extinction events, and unknown speciation mode. The only instance in which phylogenetic ages correctly predict species age is under the assumption of a bifurcating speciation process in the absence of extinction and with all living species included in the phylogenetic tree. While the prevalence of true speciation modes remains difficult to access (Bapst & Hopkins, 2017; Silvestro et al., 2018; Wagner et al., 1995), the fossil record unequivocally shows that extinction occurs across all clades in the tree of life (Bambach 2006; Benton 2023; Pimm et al., 2014), and there is substantial evidence that many living species remain unknown to science and are therefore absent from empirical phylogenetic trees (Blackwell 2001; Mora et al., 2011). Thus, the scenario under which phylogenetic age correctly predicts species age is very unlikely.

Some authors acknowledged the problems associated with measuring species age from phylogenies (Swenson, 2019), and have proposed approaches to account for them. For example, Sonne *et al*. (2022) determined young and old Andean hummingbirds by assessing the sensitivity of their results to incomplete taxon sampling by generating 1000 trees with randomly missing species. Pie & Caron (2023) accounted for taxonomic incompleteness by pruning an additional 1 – 5% of species and evaluated if their conclusions changed and found that they did not. Yet, the magnitude of the error associated with the direct use of the length of phylogenetic branches as estimators of species ages remains under-appreciated, as shown by the many studies implementing this approach.

We showed that the largest error in estimating species ages from phylogenetic trees is linked with incomplete sampling of extant species. This is a problem that in principle can be solved by extending the scope of the sampling to include all species in the phylogenetic inference. Yet, despite the advances in the scalability of DNA sequencing, this remains impractical for large clades, including some of the best sampled ones such as vertebrate groups, in which many species still lack genetic data (Jetz & Pyron 2018; Tonini et al., 2016; Upham et al., 2019). In addition, a substantial proportion of species might be missing from the phylogenetic trees because they are still unknown to science, a problem often identified as the Linnean shortfall (Diniz Filho et al., 2023; Hortal et al., 2015). The magnitude of the Linnean shortfall is unknown, but available estimates show that it affects some clades significantly more than others (Moura & Jetz 2021; Ondo et al., 2023), with the diversity of highly diverse groups, such as bacteria, insects, and fungi, likely to be highly underestimated (Blackwell 2001; Mora et al., 2011; Wiens 2023).

Our simulations showed that under some scenarios the accurate estimation of species ages is essentially impossible from phylogenetic trees using current approaches. Under the assumption of budding speciation, the error is high even without extinction and with complete sampling, because phylogenetic trees are agnostic about parent and descendant species following a branching event (Fig. 2, 4). Phylogenetic ages are, by construction, identical for sister species, which is necessarily wrong within a budding speciation scenario. Similarly, anagenetic speciation also leads to high error, which did not vary substantially with extinction. However, anagenetic speciation might be impossible to quantify, except perhaps in high resolution fossil time series (Aze et al., 2011), resulting in a general debate on the use of the term anagenesis in evolutionary biology (Vaux et al., 2015) and biogeography (Emerson & Patiño, 2018; Meiri et al., 2018). Thus, species age is unidentifiable under the assumption of speciation modes that deviate from a strictly bifurcating scenario.

The lowest error in species age estimation was observed under scenarios of bifurcating speciation. This is the implicit assumption of most studies using approximations of species ages (Alzate et al., 2023; Freer et al., 2022; Kennedy et al., 2022), even though it is at odds with the assumption of all birth-death models used in the molecular clock analyses that estimate the phylogenetic trees in the first place (Gernhard, 2008; Nee et al., 1994; Stadler, 2013). Despite the lower error, our simulations showed that both extinction and missing lineages can lead to a substantial decrease in accuracy (Fig. 4-5, SM3-4) that can even lead to qualitative misinterpretations of general patterns such as age-dependent extinction risks (Fig. 7, SM6). Given the large inaccuracy in species age, the question is whether this affects the inferences made from the relationship between species age and eco-evolutionary variables, such as extinction risk, range size, or environmental variables (Gaston & Blackburn 1997, Johnson *et al*. 2002, Tanentzap *et al*. 2015, Pie & Caron 2023).

We found our probabilistic approach to efficiently reduce the biases associated with incomplete sampling and extinction. It substantially improved the accuracy of the estimation of species ages, leading to much lower error rates even under scenarios of high extinction and high percentage of missing extant species (Fig. 6). While the error in species age estimates increased with increasing extinction, we observed the opposite pattern with our probabilistic estimates of species age under incomplete taxon sampling (Fig. SM5). This pattern is likely due to the fact that higher extinction rates set a stronger constraint on the plausible range of species age. For instance, while with a zero extinction rate species are expected to live forever (and therefore there is no *a priori* expectation on their age) a rate of e.g. 0.5 would result in an expected mean species longevity of 2 myr, thus constraining the plausible age of living species in incomplete phylogenies. The performance of our estimator is however contingent on the ability of birth-death models to correctly estimate speciation and extinction rates from phylogenies of extant species. The accuracy of these methods has been shown to be high under several simulation settings (Silvestro et al., 2011; Stadler 2011), but their accuracy has been questioned for empirical datasets and under complex models with rate variation (Louca & Pennell 2020; Rabosky 2010). One commonly observed pattern is the estimation of 0-extinction rate from empirical phylogenies (Louca & Pennell 2021). This would make corrected ages identical to the phylogenetic ages in completely sampled phylogenies, while species ages estimated from incomplete phylogenies without extinction increased in accuracy thanks to specifying the sampling fraction in our probabilistic function. Yet, the fossil record decisively shows that extinction and speciation rates vary within the same order of magnitude across virtually all clades (Parry 2021). The robustness of estimated extinction rates can be increased through the use of fossil data in the analyses (Heath et al., 2014; Silvestro et al., 2018; Warnock et al., 2020). Alternatively, our probabilistic approach could be applied across a range of plausible values of speciation rates, extinction rates, and fractions of incomplete sampling as a way to evaluate the robustness of conclusions drawn from the patterns of estimated species ages.

## Conclusion

This study quantifies the deviations between true species age and phylogenetic age due to incomplete taxon sampling, extinction, and unknown speciation modes. We found that phylogenetic age is a good proxy of species age only in a rather unlikely case in which 1) all species in a clade are known to science and included in the phylogenetic tree, 2) speciation occurs as a strictly bifurcating process, and 3) there is no or low extinction. Using simulations, we identified that incomplete taxon sampling and budding and anagenetic speciation modes cause the highest mismatch between phylogenetic age and true species age and can only be accounted for with additional information, for instance, from fossil data. In contrast, under a scenario of bifurcating speciation, we proposed a probabilistic approach based on the properties of the birth-death process that can drastically improve the accuracy of the estimated species ages by reducing the bias linked with extinction and with incomplete taxon sampling. We note that, even in this case, the robustness of such estimates will be contingent on the accuracy of the estimated speciation and extinction rates, the quantification of missing species and, of course, of the underlying phylogenetic tree and dating of the branching times. In light of our findings, we caution against the direct use of branch lengths as proxies for species ages. We recommend applying our probabilistic approach to correct species ages, exploring different values of diversification rates and incomplete sampling fractions. We hope our results will stimulate a discussion about the use of phylogenetic trees in inferring species age and lead to a critical evaluation of the robustness of inferences linking species age with traits, ecological variables, and extinction risks.

## Data and code availability

All the codes utilized in this study are available as Supplementary materials and in this repository: https://github.com/caldecid/Challenges_species_age.git

## Acknowledgements

CC received a Ph.D. scholarship from CAPES (88887.814725/2023-00) and an abroad internship CAPES-Print scholarship (88887.682496/2022-00). D.S. and TH received funding from the Swiss National Science Foundation (PCEFP3_187012). D.S. received funding the Swedish Research Council (VR: 2019-04739), and the Swedish Foundation for Strategic Environmental Research MISTRA within the framework of the research programme BIOPATH (F 2022/1448). JDC was supported by the Swiss National Science Foundation grant TMPFP2_209818.

## References

Alroy, J., Marshall, C. R., Bambach, R. K., Bezusko, K., Foote, M., Fürsich, et al., (2001). Effects of sampling standardization on estimates of phanerozoic marine diversification. Proceedings of the National Academy of Sciences of the United States of America, 98(11), 6261–6266. doi: 10.1073/pnas.111144698

Alzate, A., Rozzi, R., Velasco, J. A., Robertson, D. R., Zizka, A., Tobias, J. A., … & Onstein, R. E. (2023). The evolutionary age-range size relationship is modulated by insularity and dispersal in plants and animals. bioRxiv, 2023–11.doi: 10.1101/2023.11.11.566377

Aze, T., Ezard, T. H. G., Purvis, A., Coxall, H. K., Stewart, D. R. M., Wade, B. S., & Pearson, P. N. (2011). A phylogeny of Cenozoic macroperforate planktonic foraminifera from fossil data. Biological Reviews, 86(4), 900–927. doi: 10.1111/j.1469-185X.2011.00178.x

Balmford, A. (1996). Extinction filters and current resilience: The significance of past selection pressures for conservation biology. Trends in Ecology and Evolution. doi: 10.1016/0169-5347(96)10026-4

Bapst, D. W., & Hopkins, M. J. (2017). Comparing cal3 and other a posteriori time-scaling approaches in a case study with the pterocephaliid trilobites. Paleobiology, 43(1), 49–67. doi: 10.1017/pab.2016.34

Barido-Sottani, J., Pett, W., O’Reilly, J. E., & Warnock, R. C. M. (2019). FossilSim: An R package for simulating fossil occurrence data under mechanistic models of preservation and recovery. Methods in Ecology and Evolution, 10(6), 835–840. doi: 10.1111/2041-210X.13170

Barraclough, T. G., & Vogler, A. P. (2000). Detecting the geographical pattern of speciation from species-level phylogenies. The American Naturalist, 155(4), 419–434. doi: 10.1086/303332

Barraclough, T. G., Vogler, A. P., & Harvey, P. H. (1998). Revealing the factors that promote speciation. Philosophical Transactions of the Royal Society B: Biological Sciences, 353(1366), 241–249. doi: 10.1098/rstb.1998.0206

Bambach, R. K. (2006). Phanerozoic biodiversity mass extinctions. Annual Review of Earth and Planetary Sciences, 34, 127–155. doi: 10.1146/annurev.earth.33.092203.122654

Baum, D. A., Smith, S. D. W., & Donovan, S. S. S. (2005). The tree-thinking challenge. Science, 310(5750), 979–980. doi: 10.1126/science.1117727

Beaulieu, J. M., & O’Meara, B. C. (2016). Detecting hidden diversification shifts in models of trait-dependent speciation and extinction. Systematic Biology, 65(4), 583–601. doi: 10.1093/sysbio/syw022

Benton, M. J. (2013). Origins of biodiversity. Palaeontology. doi: 10.1111/pala.12012

Benton, M. J. (2016). Origins of Biodiversity. PLoS Biology, 14(11), 1–7. doi: 10.1371/journal.pbio.2000724

Benton, M. J. (2023). Extinctions: How life survives, adapts and evolves. Thames & Hudson.

Blackwell, M. (2011). The Fungi: 1, 2, 3… 5.1 million species?. American journal of botany, 98(3), 426–438. doi: 10.3732/ajb.1000298

Brée, B., Condamine, F. L., & Guinot, G. (2022). Combining palaeontological and neontological data shows a delayed diversification burst of carcharhiniform sharks likely mediated by environmental change. Scientific Reports, 12(1), 21906. doi: 10.1038/s41598-022-26010-7

Caetano, D. S., & Quental, T. B. (2022). How important is budding speciaiton for comparative studies? BioRxiv. doi: 10.1101/2022.05.24.493296

Callaway, E. (2017). Oldest *Homo sapiens* fossil claim rewrites our species’ history. Nature, 546, 289–293. doi: 10.1038/nature.2017.22114

Carrillo, J. D., Faurby, S., Silvestro, D., Zizka, A., Jaramillo, C., Bacon, C. D., & Antonelli, A. (2020). Disproportionate extinction of South American mammals drove the asymmetry of the Great American Biotic Interchange. Proceedings of the National Academy of Sciences of the United States of America, 117(42), 26281–26287. doi: 10.1073/pnas.2009397117

Carrillo, J. D., Forasiepi, A., Jaramillo, C., Sánchez-Villagra, M. R., & Richardson, J. E. (2015). Neotropical mammal diversity and the Great American Biotic Interchange: spatial and temporal variation in South America’s fossil record. Frontiers in genetics, 5, 451. doi: 10.3389/fgene.2014.00451

Chang, J., Rabosky, D. L., & Alfaro, M. E. (2020). Estimating diversification rates on incompletely sampled phylogenies: theoretical concerns and practical solutions. Systematic Biology, 69(3), 602–611. doi: 10.1093/sysbio/syz081

Davies, T. J., Smith, G. F., Bellstedt, D. U., Boatwright, J. S., Bytebier, B., Cowling, R. M., et al., (2011). Extinction risk and diversification are linked in a plant biodiversity hotspot. PLoS Biology. 9(5), e1000620. doi: 10.1371/journal.pbio.1000620

Diniz Filho, J. A. F., Jardim, L., Guedes, J. J., Meyer, L., Stropp, J., Frateles, L. E. F., … & Hortal, J. (2023). Macroecological links between the Linnean, Wallacean, and Darwinian shortfalls. Frontiers of Biogeography, 15(2). doi: 10.21425/F5FBG59566

Eldredge, N., Thompson, J. N., Brakefield, P. M., Gavrilets, S., Jablonski, D., Jackson, J. B. C., et al., (2005). The dynamics of evolutionary stasis. Paleobiology 31(S2), 133–145. doi: 10.1666/0094-8373(2005)031

Emerson, B. C., & Patiño, J. (2018). Anagenesis, cladogenesis, and speciation on islands. Trends in Ecology and Evolution, 33(7), 488–491. doi: 10.1016/j.tree.2018.04.006

Fitzpatrick, B. M., & Turelli, M. (2006). The geography of mammalian speciation: mixed signals from phylogenies and range maps. Evolution, 60(3), 601–615. doi: 10.1111/j.0014-3820.2006.tb01140.x

Fjeldså, J., Bowie, R. C. K., & Rahbek, C. (2012). The role of mountain ranges in the diversification of birds. Annual Review of Ecology, Evolution, and Systematics, 43, 249–265. doi: 10.1146/annurev-ecolsys-102710-145113

Foote, M. (1996). On the probability of ancestors in the fossil record. Paleobiology, 22(2), 141–151. doi: 10.1017/S0094837300016146

Foote, M., & Raup, D. M. (1996). Fossil preservation and the stratigraphic ranges of taxa. Paleobiology, 22(2), 121–140. doi: 10.1017/S0094837300016134

Freer, J. J., Collins, R. A., Tarling, G. A., Collins, M. A., Partridge, J. C., & Genner, M. J. (2022). Global phylogeography of hyperdiverse lanternfishes indicates sympatric speciation in the deep sea. Global Ecology and Biogeography, 31(11), 2353–2367. doi: 10.1111/geb.13586

Gaston, K. J., & Blackburn, T. M. (1997). Evolutionary age and risk of extinction in the global avifauna. Evolutionary Ecology, 11, 557–565. doi: 10.1007/s10682-997-1511-4

Gernhard, T. (2008). The conditioned reconstructed process. Journal of theoretical biology, 253(4), 769–778. doi: 10.1016/j.jtbi.2008.04.005

Harvey, P. H., May, R. M., & Nee, S. (1994). Phylogenies without fossils. Evolution, 48(3), 523–529. doi: 10.1111/j.1558-5646.1994.tb01341.x

Heath, T. A., Hedtke, S. M., & Hillis, D. M. (2008). Taxon sampling and the accuracy of phylogenetic analyses. Journal of Systematics and Evolution, 46(3), 239–257. doi: 10.3724/SP.J.1002.2008.08016

Heath, T. A., Huelsenbeck, J. P., & Stadler, T. (2014). The fossilized birth–death process for coherent calibration of divergence-time estimates. Proceedings of the National Academy of Sciences, 111(29), E2957–E2966. doi: 10.1073/pnas.1319091111

Hortal, J., de Bello, F., Diniz-Filho, J. A. F., Lewinsohn, T. M., Lobo, J. M., & Ladle, R. J. (2015). Seven shortfalls that beset large-scale knowledge of biodiversity. Annual Review of Ecology, Evolution, and Systematics, 46, 523–549. doi: 10.1146/annurev-ecolsys-112414-054400

IUCN. (2016). IUCN Red List of Threatened Species. Version 2016-2. Fourth Quarter.

Jetz, W., & Pyron, R. A. (2018). The interplay of past diversification and evolutionary isolation with present imperilment across the amphibian tree of life. Nature ecology & evolution, 2(5), 850–858. doi: 10.1038/s41559-018-0515-5

Johnson, C. N., Delean, S., & Balmford, A. (2002). Phylogeny and the selectivity of extinction in Australian marsupials. 135–142. doi: 10.1017/S1367943002002196

Kendall, M. G. (1946). The advanced theory of statistics (2nd ed.). Charles Griffin and Co., Ltd., London.

Kennedy, J. D., Marki, P. Z., Reeve, A. H., Blom, M. P., Prawiradilaga, D. M., Haryoko, T., … & Jønsson, K. A. (2022). Diversification and community assembly of the world’s largest tropical island. Global Ecology and Biogeography, 31(6), 1078–1089. doi: 10.1111/geb.13484

López-Martínez, A. M., Schonenberger, J., von Balthazar, M., González-Martínez, C. A., Ramírez-Barahona, S., Sauquet, H., & Magallón, S. (2023). Integrating fossil flowers into the Angiosperm phylogeny using molecular and morphological evidence. Systematic Biology, syad017. doi: 10.1093/sysbio/syad017

Losos, J. B., & Glor, R. E. (2003). Phylogenetic comparative methods and the geography of speciation. Trends in Ecology and Evolution, 18(5), 220–227. doi: 10.1016/S0169-5347(03)00037-5

Louca, S., & Pennell, M. W. (2020). Extant timetrees are consistent with a myriad of diversification histories. Nature, 580(7804), 502–505. doi: 10.1038/s41586-020-2176-1

Louca, S., & Pennell, M. W. (2021). Why extinction estimates from extant phylogenies are so often zero. Current Biology, 31(14), 3168–3173. doi:10.1016/j.cub.2021.04.066

Meiri, S., Raia, P., & Santos, A. M. (2018). Anagenesis and cladogenesis are useful island biogeography terms. Trends in Ecology & Evolution, 33(12), 895–896. doi: 10.1016/j.tree.2018.09.005

Mora, C., Tittensor, D. P., Adl, S., Simpson, A. G., & Worm, B. (2011). How many species are there on Earth and in the ocean?. PLoS biology, 9(8), e1001127. doi: 10.1371/journal.pbio.1001127

Moura, M. R., & Jetz, W. (2021). Shortfalls and opportunities in terrestrial vertebrate species discovery. Nature Ecology & Evolution, 5(5), 631–639. doi: 10.1038/s41559-021-01411-5

Mynard, P., Algar, A., Lancaster, L., Bocedi, G., Fahri, F., Gubry-Rangin, C., et al., (2023). Impact of phylogenetic tree completeness and misspecification of sampling fractions on trait dependent diversification models. Systematic Biology, 72(1), 106–119. doi: 10.1093/sysbio/syad001

Nee, S., May, R. M., & Harvey, P. H. (1994). The reconstructed evolutionary process. Philosophical Transactions of the Royal Society of London. Series B: Biological Sciences, 344(1309), 305–311. doi: 10.1098/rstb.1994.0068

Nee, S., & May, R. M. (1997). Extinction and the loss of evolutionary history. Science, 278(5338), 692–694. doi: 10.1126/science.278.5338.692

Ondo, I., Dhanjal-Adams, K., Pironon, S., Silvestro, D., Deklerck, V., Grace, O., … & Antonelli, A. (2023). Plant diversity darkspots for global collection priorities. bioRxiv, 2023–09.doi: 10.1101/2023.09.12.557387

Parry, L. A. (2021). Evolution: no extinction? No way!. Current Biology, 31(14), R907–R909. doi: 10.1016/j.cub.2021.05.044

Pearson, P. N. (1995). Investigating age-dependency of species extinction rates using dynamic survivorship analysis. Historical Biology, 10(2), 119–136. doi: 10.1080/10292389509380516

Pie, M. R., & Caron, F. (2023). Substantial variation in species ages among vertebrate clades. BioRxiv, 06. doi: 10.1101/2023.06.08.544238

Pimm, S. L., Jenkins, C. N., Abell, R., Brooks, T. M., Gittleman, J. L., Joppa, L. N., … Sexton, J. O. (2014). The biodiversity of species and their rates of extinction, distribution, and protection. Science, 344(6187). doi: 10.1126/science.1246752

R Core Team. (2019). R: A language and environment for statistical computing. Vienna, Austria: R Foundation for Statistical Computing. Retrieved from https://www.r-project.org/

Rabosky, D. L. (2010). Extinction rates should not be estimated from molecular phylogenies. Evolution, 64(6), 1816–1824. doi: 10.1111/j.1558-5646.2009.00926.x

Rivas-Gonzáles, I., Rousselle, M., Li, F., Zhou, L., Dutheil, J., Munch, K., et al., (2023). Pervasive incomplete lineage sorting illuminates speciation and selection in primates. Science, 380(6648), eabn4409. doi: 10.1126/science.abn4409

Roopnarine, P. D., Byars, G., Fitzgerald, P., Paleobiology, S., & Winter, N. (1999). Anagenetic evolution, stratophenetic patterns, and random walk models. Paleobiology, 25(1), 41–57. doi: 10.1666/0094-8373(1999)0252.3.CO;2

Rosenblum, E. B., Sarver, B. A. J., Brown, J. W., Des Roches, S., Hardwick, K. M., Hether, T. D., et al., (2012). Goldilocks meets Santa Rosalia: an ephemeral speciation model explains patterns of diversification across time scales. Evolutionary Biology, 39(2), 255–261. doi: 10.1007/s11692-012-9171-x

Silvestro, D., Schnitzler, J., & Zizka, G. (2011). A Bayesian framework to estimate diversification rates and their variation through time and space. BMC evolutionary biology, 11, 1–15. doi: 10.1186/1471-2148-11-311

Silvestro, D., Salamin, N., & Schnitzler, J. (2014). PyRate: a new program to estimate speciation and extinction rates from incomplete fossil data. Methods in Ecology and Evolution2, 5(10), 1126–1131. doi: 10.1111/2041-210X.12263

Silvestro, D., Castiglione, S., Mondanaro, A., Serio, C., Melchionna, M., Piras, P., et al., (2020). A 450 million years long latitudinal gradient in age-dependent extinction. Ecology Letters, 23(3), 439–446. doi: 10.1111/ele.13441

Silvestro, D., Salamin, N., Antonelli, A., & Meyer, X. (2019). Improved estimation of macroevolutionary rates from fossil data using a Bayesian framework. Paleobiology, 45(4), 546–570. doi: 10.1017/pab.2019.23

Silvestro, D., Warnock, R. C. M., Gavryushkina, A., & Stadler, T. (2018). Closing the gap between palaeontological and neontological speciation and extinction rate estimates. Nature Communications, 9(1). doi: 10.1038/s41467-018-07622-y

Simpson, George Gaylord. (1984). Tempo and mode in evolution. Columbia University Press.

Skeels, A., & Cardillo, M. (2019). Reconstructing the geography of speciation from contemporary biodiversity data. American Naturalist, 193(2), 240–255. doi: 10.1086/701125

Slater, G. J., & Harmon, L. J. (2013). Unifying fossils and phylogenies for comparative analyses of diversification and trait evolution. Methods in Ecology and Evolution, 4(8), 699–702. doi: 10.1111/2041-210X.12091

Sonne, J., Dalsgaard, B., Borregaard, M. K., Kennedy, J., Fjeldså, J., & Rahbek, C. (2022). Biodiversity cradles and museums segregating within hotspots of endemism. Proceedings of the Royal Society B: Biological Sciences, 289(1981), 20221102. doi: 10.1098/rspb.2022.1102

Stadler, T. (2011). Mammalian phylogeny reveals recent diversification rate shifts. Proceedings of the National Academy of Sciences, 108(15), 6187–6192. doi: 10.1073/pnas.1016876108

Stadler, T. (2013). Recovering speciation and extinction dynamics based on phylogenies. Journal of evolutionary biology, 26(6), 1203–1219. doi: 10.1111/jeb.12139

Stadler, T., Gavryushkina, A., Warnock, R. C., Drummond, A. J., & Heath, T. A. (2018). The fossilized birth-death model for the analysis of stratigraphic range data under different speciation modes. Journal of theoretical biology, 447, 41–55. doi: 10.1016/j.jtbi.2018.03.005

Stadler T (2019). TreeSim: Simulating Phylogenetic Trees. R package version 2. 4, https://CRAN.R-project.org/package=TreeSim

Swenson, N. G. (2019). Phylogenetic ecology: A history, critique, and remodeling. University of Chicago Press.

Tanentzap, A. J., Brandt, A. J., Smissen, R. D., Heenan, P. B., Fukami, T., & Lee, W. G. (2015). When do plant radiations influence community assembly? The importance of historical contingency in the race for niche space. New Phytologist, 207(2), 468–479. doi: 10.1111/nph.13362

Tanentzap, A. J., Igea, J., Johnston, M. G., & Larcombe, M. J. (2020). Does evolutionary history correlate with contemporary extinction risk by influencing range size dynamics? American Naturalist, 195(3), 569–576. doi: 10.1086/707207

Thomas, G. H., Hartmann, K., Jetz, W., Joy, J. B., Mimoto, A., & Mooers, A. O. (2013). PASTIS: An R package to facilitate phylogenetic assembly with soft taxonomic inferences. Methods in Ecology and Evolution, 4(11), 1011–1017. doi: 10.1111/2041-210X.12117

Tonini, J. F. R., Beard, K. H., Ferreira, R. B., Jetz, W., & Pyron, R. A. (2016). Fully-sampled phylogenies of squamates reveal evolutionary patterns in threat status. Biological Conservation, 204, 23–31. doi: 10.1016/j.biocon.2016.03.039

Upham, N. S., Esselstyn, J. A., & Jetz, W. (2019). Inferring the mammal tree: Species-level sets of phylogenies for questions in ecology, evolution, and conservation. In PLoS Biology, 17(12), e3000494. doi: 10.1371/journal.pbio.3000494

Van Valen, L. (1973). A new evolutionary law. Evolutionary Theory. 1, 1–30.

Vaux, F., Trewick, S. A., & Morgan-Richards, M. (2016). Lineages, splits and divergence challenge whether the terms anagenesis and cladogenesis are necessary. Biological Journal of the Linnean Society, 117(2), 165–176. doi: 10.1111/bij.12665

Verde Arregoitia, L. D., Blomberg, S. P., & Fisher, D. O. (2013). Phylogenetic correlates of extinction risk in mammals: Species in older lineages are not at greater risk. Proceedings of the Royal Society B: Biological Sciences 280(1765), 20131092. doi: 10.1098/rspb.2013.1092

Wagner, P. J., Erwin, D. H., & Anstey, R. L. (1995). Phylogenetic patterns as tests of speciation models. In New approaches to speciaiton in the fossil record (pp. 87–122). New York: Columbia University Press.

Warnock, R. C., Heath, T. A., & Stadler, T. (2020). Assessing the impact of incomplete species sampling on estimates of speciation and extinction rates. Paleobiology, 46(2), 137–157. doi: 10.1017/pab.2020.12

Wiens, J. J. (2023). How many species are there on Earth? Progress and problems. PLoS biology, 21(11), e3002388. doi: 10.1371/journal.pbio.3002388

Yang, Z., & Rannala, B. (1997). Bayesian phylogenetic inference using DNA sequences: a Markov Chain Monte Carlo method. Molecular biology and evolution, 14(7), 717–724. doi: 10.1093/oxfordjournals.molbev.a025811

