## Supplement for "Challenges in estimating species age from phylogenetic trees"

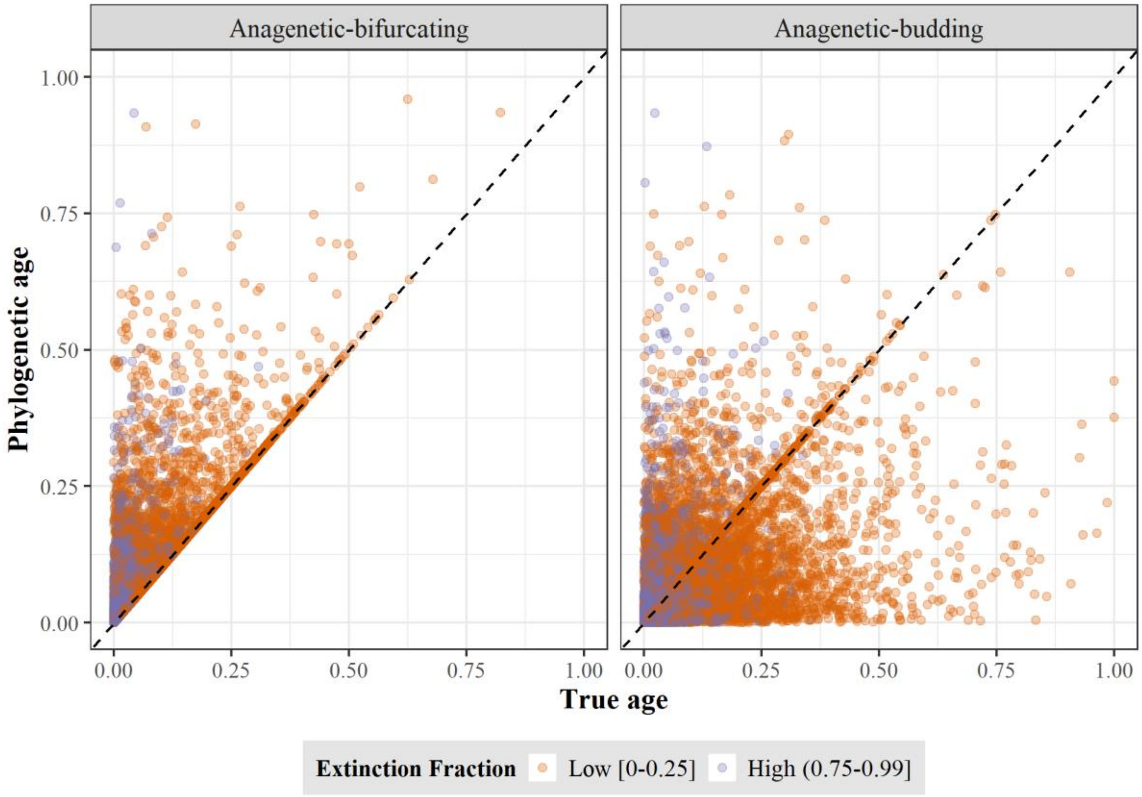


**Fig. SM1.** True age versus phylogenetic age at low and high extinction fraction for Anagenetic-bifurcating (left) and Anagenetic-budding (right) speciation. Each point represents a species. True and phylogenetic ages are scaled to the root age of the correspondent phylogenetic tree.

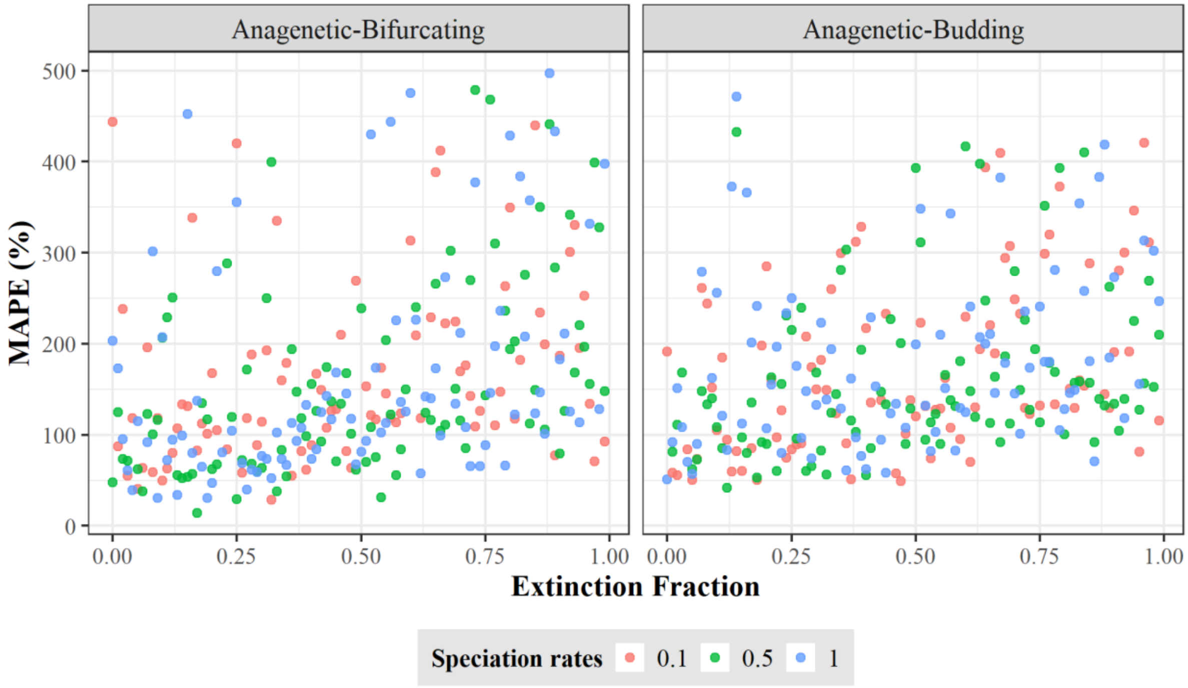


**Fig. SM2.** Error in equating phylogenetic age with speciation age. The error was quantified as mean absolute percentage error (MAPE) between the true and phylogenetic ages across all species for each tree simulated under Anagenetic-bifurcating (left) and Anagenetic-budding speciation (right). Each dot represents one replicate of the 300 trees for each speciation mode using different rates of speciation and extinction fraction.

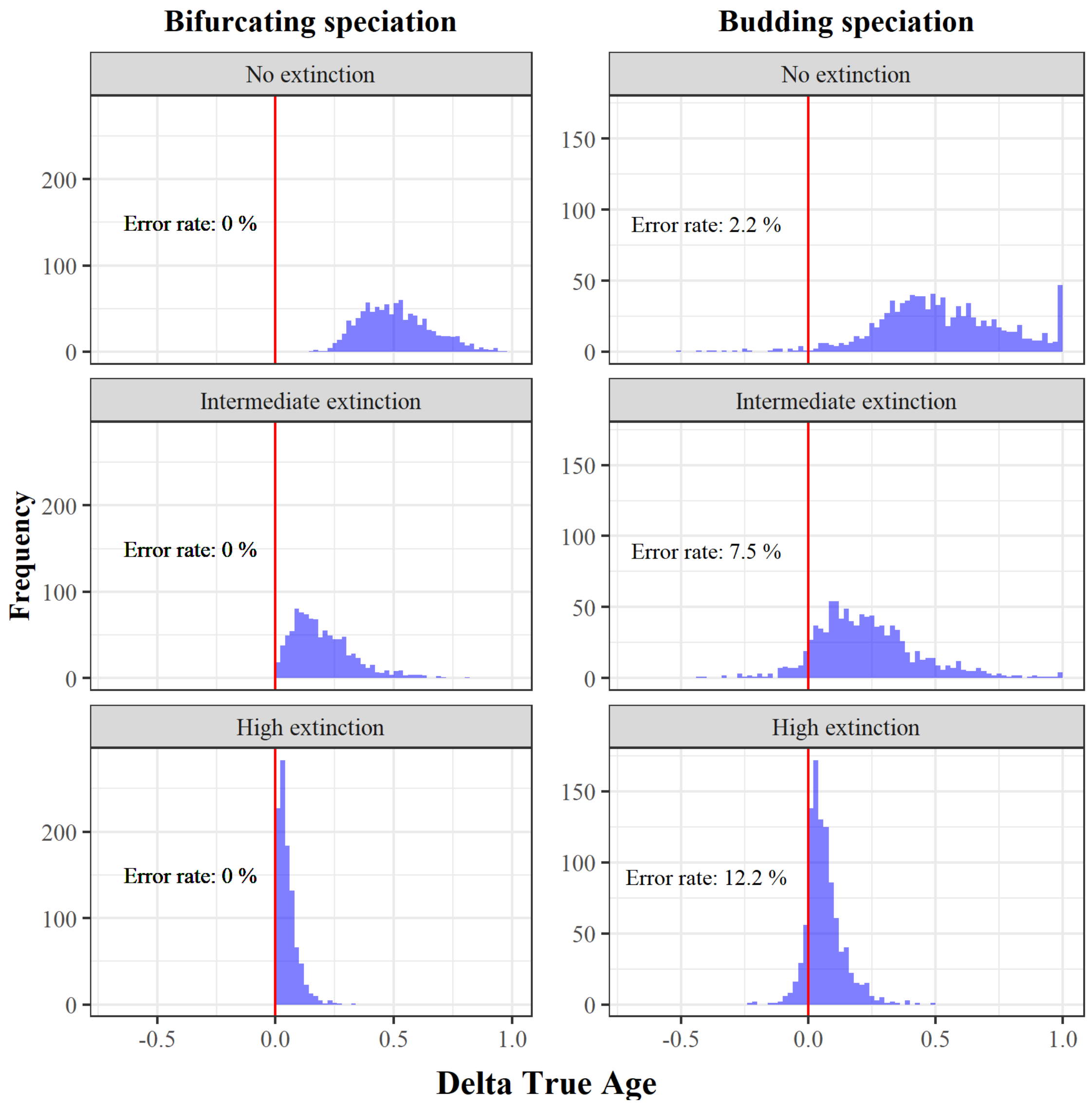


**Fig. SM3.** Error in estimating the relative age of species. For each of the 1000 simulations under bifurcating (left) and budding (right) speciation, combined with three different extinction levels, we selected the oldest and youngest species according to the phylogenetic ages, and calculated the difference in their true ages (ΔTrue age). A ΔTrue age smaller than 0 indicates that the phylogenetic oldest species was estimated to be in fact younger than the phylogenetic youngest species, and therefore, the comparison of phylogenetic ages is qualitatively wrong.

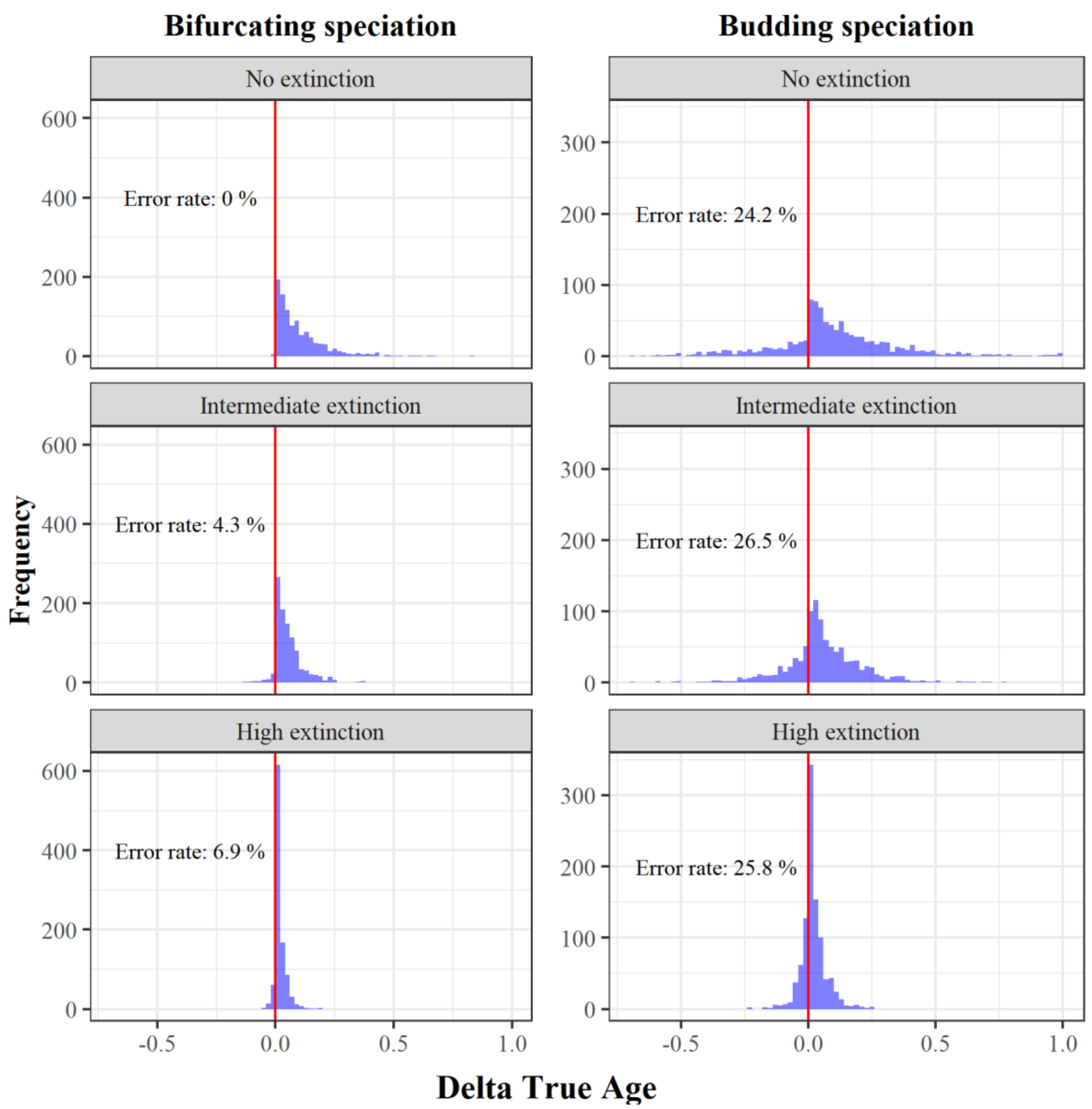


**Fig. SM4.** Risk to confuse older with younger random species. For each of the 1000 simulations under bifurcating (left) and budding (right) speciation, combined with three different extinction levels, we selected two random species and defined which was the older and younger according to the phylogenetic ages, and calculated the difference in their true ages (ΔTrue age). A ΔTrue age smaller than 0 indicates that the phylogenetic older species was in fact younger than the phylogenetic younger species, and therefore, the comparison of phylogenetic ages is qualitatively wrong.

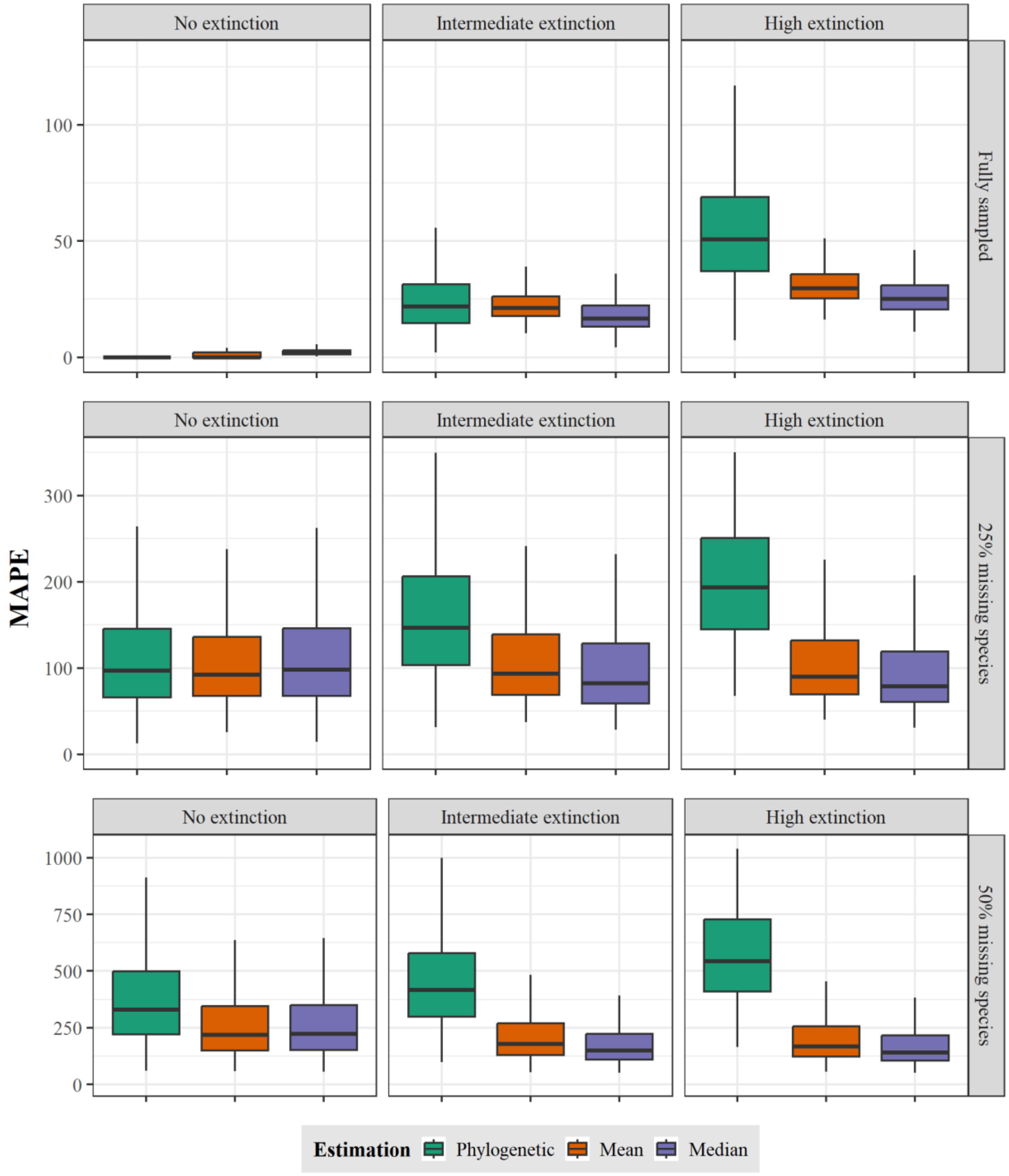


**Fig. SM5**. Performance of a probabilistic age estimator. Error in equating the phylogenetic age and the probability estimator point estimates (mean and median) with the true species age for three extinction scenarios (no extinction, intermediate, and high; from left to right) and three sampling scenarios (fully sampled, 25%, and 50% missing species; from up to down). The error was quantified as mean absolute percentage error (MAPE) between the true and point estimates or phylogenetic ages across 100 species for each of 1000 trees for each extinction scenario simulated under bifurcating speciation.

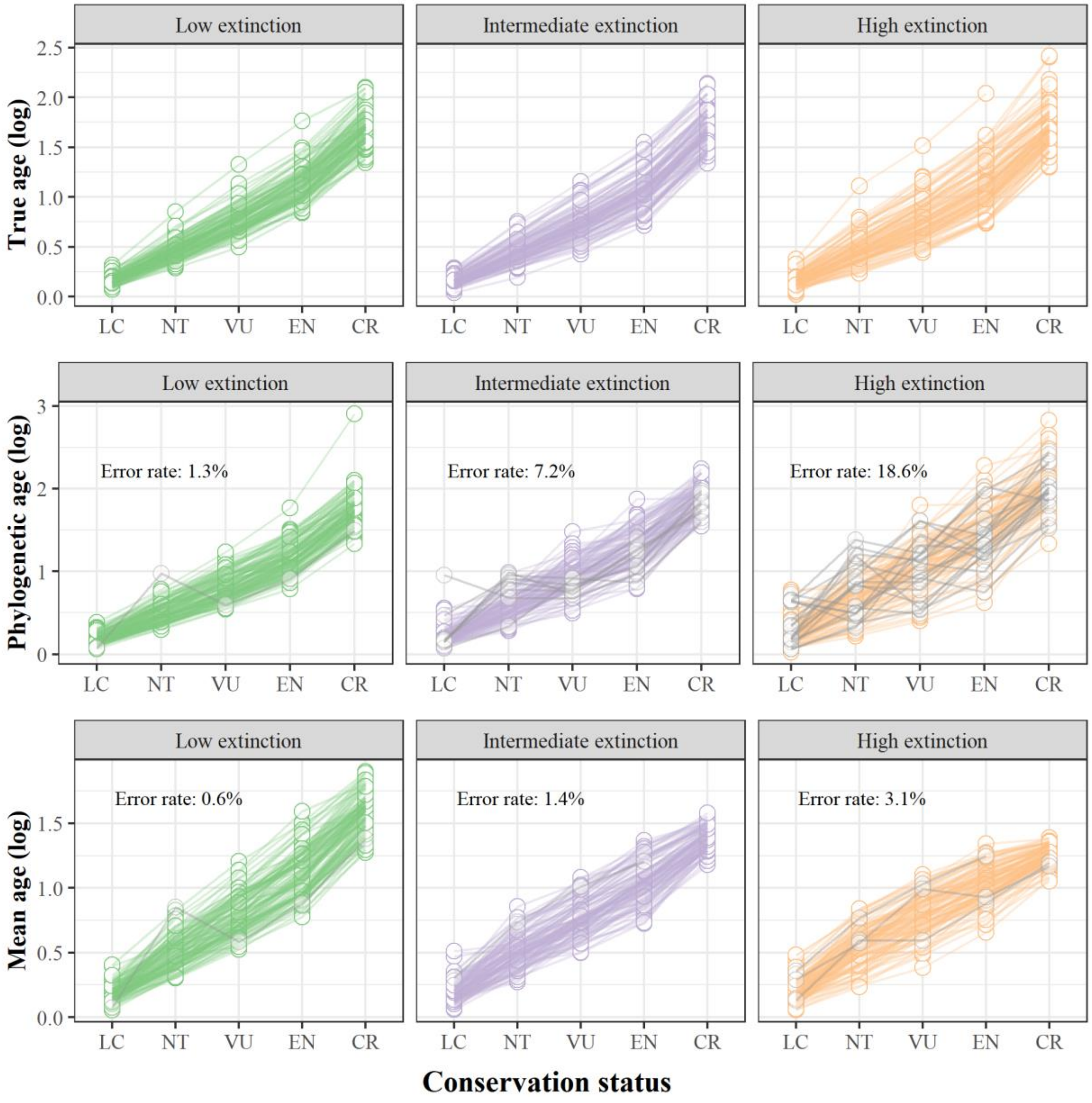


**Fig. SM6.** Power to recover an age extinction-risk relationship. Simulated species ages under three extinction scenarios (low, intermediate, and high; from left to right) and assuming bifurcating speciation and fully sampled trees were binned into conservation status categories, which represents an increase in extinction risk by age (LC = Least Concern; NT = Near Threatened; VU = Vulnerable; EN = Endangered; CR = Critically Endangered). We used the phylogenetic age and the mean age obtained from our probabilistic corrective function to calculate the mean age per conservation status category and assess if every mean age increases in comparison with the previous category with lower extinction risk. The error rate is the percentage across all 1000 phylogenies where the relationship between the mean ages and the conservation status categories is not always increasing (shown by grey lines).
